# Cancer-Testis Antigens as Clinical and Prognostic Biomarkers in Gastric Adenocarcinoma: Integration of Differential Expression, Clinical Associations, Survival, and Co-Expression Networks

**DOI:** 10.1101/2025.09.17.676427

**Authors:** Jade Dias Valente, Livia Érika Carlos Marques, Ronald Matheus da Silva Mourão, Bianca de Fátima dos Reis Rodrigues, Louise Sousa de Souza, Jéssica Manoelli Costa da Silva, Samia Demachki, Geraldo Ishak, Fabiano Cordeiro Moreira, Samir Mansour Moraes Casseb, Rommel Mario Rodriguez Burbano, Paulo Pimentel de Assumpção

## Abstract

Gastric adenocarcinoma (GAC) remains one of the leading causes of cancer-related mortality, characterized by marked molecular and clinical heterogeneity, which underscores the need for robust biomarkers. In this study, we aimed to evaluate the role of cancer-testis antigens (CTAs) in GAC through integrated analyses of differential expression, clinical associations, survival impact, and gene co-expression networks. The cohort included 156 patients with complete clinical data, comprising a total of 362 tissue samples (tumor, peritumoral, and metaplastic). We identified 541 differentially expressed genes, of which 49 corresponded to previously described CTAs. Between GAC and peritumoral tissues, 14 CTAs exhibited significant differential expression, with MAGEA3, MAGEA6, GOLGA6L1, and MAGEA2 among the most highly expressed in tumors. Relevant associations were observed between the expression of MAGEA3, MAGEA6, POTEF, and CTCFL with TNM staging, as well as IGF2BP1, DAZ1/3/4, YBX2, and PCDHA4 in relation to variables such as tumor depth, metastasis, and TCGA subtypes. Survival analysis demonstrated that high expression of IGF2BP1, CTCFL, CT45A5, and LIN28B was strongly associated with worse prognosis (HR > 2.7), whereas SYCE1L, PIK3R3, and ZNF683 showed a protective effect. The co-expression network revealed five main clusters, highlighting germline-related modules (MAGEA, DAZ, CSAG), adhesion and transcriptional regulation (YBX2, POTEF, PCDHA, TAF1L), and immune-related genes (IRAK3, CXCR1, PIK3R3), evidencing functional integration between proliferation, adhesion, and immune microenvironment modulation. Collectively, these findings reinforce CTAs as potential clinical and prognostic biomarkers in GAC, with direct implications for risk stratification and the development of personalized therapeutic strategies.

## Introduction

Gastric cancer (GC) remains a major global health challenge, ranking as the fourth leading cause of cancer-related mortality worldwide [1]. Most patients have a poor prognosis, with five-year survival rates ranging from 20% to 30% in most regions [2], primarily due to late-stage diagnosis [3].

The standard approach to GC is based on surgical intervention, often combined with neoadjuvant chemotherapy and radiotherapy or immunotherapy [4]. In recent decades, immunotherapy has emerged as an alternative option for cancers unresponsive to conventional treatments, representing a promising strategy to improve the prognosis of patients with GC. Consequently, increasing research efforts have focused on the development of more selective targeted therapies and immunotherapies, as well as on the identification of novel predictive biomarkers and therapeutic targets [5].

In this context, cancer-testis antigens (CTAs) are proteins encoded by a family of genes with an expression profile restricted to germline and trophoblastic cells, but also highly expressed in different types of cancer [6]. CTAs have been implicated in promoting tumorigenesis, therapy resistance, and metastatic progression [7,8]. They are also involved in multiple cancer-related cellular pathways and contribute to nearly all cancer hallmarks, from the original to the emerging ones [9].

Moreover, CTAs exhibit unique characteristics that position them as attractive targets for immune-directed cancer therapies [10]. The testis and placenta, the tissues in which they are expressed, constitute immune-privileged sites due to the presence of the blood–testis and blood–placenta barriers, which limit immune system access and prevent the establishment of long-term immune tolerance to these antigens [11]. Consequently, CTAs aberrantly expressed in other tissues are recognized as “non-self” and elicit cytotoxic T lymphocyte (CTL)-mediated responses, triggering antitumor immunity [12]. Accordingly, CTAs are currently being evaluated in a wide range of clinical trials [13–15].

In 2009, more than 200 CTA-coding genes were cataloged in a dedicated database (http://www.cta.lncc.br/) [16]. Since then, new technologies have enabled a significant expansion in the number of identified CTAs [17], although these have not been incorporated into the database. Consequently, the lack of a comprehensive and updated registry of genes classified as CTAs means that most studies investigating CTA gene expression patterns in cancer rely on outdated lists [18, 19], which limits both the interpretation and reproducibility of findings.

Advances in next-generation sequencing (NGS) technologies enable transcriptomic data analyses, allowing the identification of specific expression patterns and the detection of differentially expressed genes (DEGs), thereby providing valuable insights for the discovery of potential biomarkers and novel therapeutic targets [20,21]. The integration of new data with existing knowledge may offer additional benefits for gastric cancer immunotherapy in the coming years [22].

In this study, we investigated the transcriptional profile of CTAs by analyzing their differential expression in gastric adenocarcinoma (GAC), peritumoral tissue (PTT), and metaplastic tissue (MP). The identified DEGs were correlated with clinical and epidemiological variables, as well as their impact on overall survival. Additionally, we constructed co-expression networks to explore the functional interactions between CTAs and partner genes. This study provides comprehensive insights into GAC and underscores the relevance of CTAs as potential biomarkers and therapeutic targets for immunotherapy.

## Methods

### Ethical Considerations and Sample Characterization

A total of 362 samples were collected, divided into three categories: 156 GAC samples, including both Lauren histological types, EBV-positive and EBV-negative cases, and *H. pylori*-positive and *H. pylori*-negative cases; 186 peritumoral tissue (PTT) samples adjacent to the tumor; and 20 metaplastic tissue (MP) samples, a precursor condition that may progress to gastric cancer. During gastric resection, 0.5 cm fragments were collected from each tumor, 0.5 cm from MP tissue, and another 0.5 cm fragment from tissue located at least 5 cm away from the macroscopic tumor margins, for cases included in the PTT analysis. Samples were stored in RNAlater and subsequently preserved at −80 °C.

The medical records of patients included in this study were reviewed to obtain clinical and epidemiological data, including sex, age, Lauren histological subtype, TCGA classification, TNM staging, tumor location, therapeutic management, and T, N, and M categories. Participants were recruited from the João de Barros Barreto University Hospital (HUJBB) of the Federal University of Pará and the Ophir Loyola Hospital (HOL). All study participants provided written informed consent, voluntarily agreed to participate, and were informed about the study objectives. This study was conducted with the approval of the Research Ethics Committee under protocol number CAAE 47580121.9.0000.5634.

### Total RNA Extraction and Sequencing

Total RNA from tissue samples was extracted using TRIzol® (Thermo Fisher Scientific), following the manufacturer’s instructions. Approximately 50–100 mg of tissue from each sample was initially homogenized, after which 1 mL of TRIzol was added to the processed tissue at room temperature. RNA integrity and concentration were assessed using the Qubit 4.0 Fluorometer and the NanoDrop ND-1000 (Thermo Fisher Scientific). Optimal criteria for total RNA integrity were defined as A260/A280 ratios between 1.8 and 2.2, A260/A230 ratios >1.8, and RNA Integrity Number (RIN) ≥5. The extracted RNA was stored at –80 °C until further use.

### Library Preparation and NGS Sequencing

Library preparation was performed using 1 μg of total RNA per sample in a final volume of 10 μL, with the TruSeq Stranded Total RNA Library Prep Kit with Ribo-Zero Gold (Illumina), following the manufacturer’s instructions. After library construction, integrity was reassessed using the 2200 TapeStation System (Agilent), and the final product showed a fragment of approximately 260 base pairs. cDNA libraries were sequenced on the Illumina NextSeq platform in paired-end mode, using the commercial NextSeq® 500 High Output Kit V2, 150 cycles (Illumina®), under the conditions specified by the manufacturer.

### Read Quality Control

Reads obtained were converted to FASTQ format using Reporter software, with all sequences and quality scores encoded in ASCII. Read quality was assessed with FastQC (v0.11.9), and adapters and low-quality reads were subsequently removed using Trimmomatic [23]. Reads were filtered and aligned to the human transcriptome reference (hg38) using Salmon (v1.5.2) [24] in two steps: quantification of mRNAs using FASTA sequences of human coding transcripts, and identification of ncRNAs using FASTA sequences of non-coding transcripts. The resulting transcript abundances were imported into R (v4.5.1; The R Project for Statistical Computing) [25] using the Tximport package [26].

### Differential Transcript Expression Analysis

The resulting count matrices were normalized, and gene-level transcript abundances were estimated using the DESeq2 package [27] in RStudio. Differential expression analysis was performed across the three sample groups, with comparisons conducted through specific contrasts between the groups of interest. Transcripts were considered differentially expressed if they met the following criteria: |log2(fold change)| (L2FC) > 2 and adjusted p-value < 0.05. Results were visualized using a volcano plot, and normalized counts were extracted for downstream analyses.

### CTA Gene Selection

In this study, a comprehensive list of 1,103 putative CTA genes, previously described by Silva et al. [17], was used. In that work, the authors identified genes with preferential or exclusive expression in the testis based on RNA sequencing (RNA-Seq) data from the Human Body Map (HBM), Genotype-Tissue Expression Project (GTEx), and Human Protein Atlas (HPA) databases. As a selection criterion, a proportional score ≥ 0.9 was applied, indicating that at least 90% of the total gene expression across the evaluated tissues originated from the testis. For subsequent analyses, this previously established CTA gene list (n = 1,103 genes) was directly adopted.

### Association of CTAs with Clinical Variables

Normalized data were used to assess the relationship between CTA gene expression in GAC and i) pathological variables, including Lauren histological subtype, TCGA classification, TNM staging, and T, N, and M categories, and ii) epidemiological characteristics, including sex and age. Associations were evaluated using DESeq2 in RStudio, with an adjusted p-value < 0.05. Additionally, non-parametric Kruskal–Wallis and Wilcoxon tests (p < 0.05) were applied as complementary approaches to validate the results obtained with DESeq2. Results were visualized using boxplots.

### Overall Survival Analysis

For overall survival (OS) analysis, multivariate Cox regression models with 95% confidence intervals were applied using the Survival package in R. P-values were obtained from both the log-rank test and the Cox model, with the corresponding hazard ratios (HRs) reported. Genes with p < 0.05 were considered statistically significant. Kaplan–Meier curves were generated using the Survminer package to visualize survival differences between high and low gene expression groups.

### Construction of Co-expression Networks

Starting from the variance-stabilizing transformation (VST)–normalized expression matrix, we defined as target genes the cancer-testis antigens (CTAs) previously identified as differentially expressed between tumor and non-tumor samples. For each CTA, we calculated the Spearman correlation (ρ) between its expression and that of all other genes, selecting the five partners with the highest |ρ| values for each target. This strategy focused the analysis on the most informative “context” of each CTA, reducing noise and multiple-testing issues while generating a biologically interpretable subnetwork.

We then constructed an undirected graph weighted by |ρ|, preserving the correlation sign as an edge attribute, and applied a |ρ| ≥ 0.30 filter to retain only relevant associations. Cluster detection was performed using the Leiden algorithm under the Constant Potts Model (CPM), prioritizing the identification of macro-modules of co-expression around CTAs, useful for generating hypotheses of gene coregulation and shared pathway involvement. The visualization layout was obtained with the layout_with_graphopt function from the igraph package (version 1.3.5). Networks were rendered with the ggraph package (version 2.2.2), and a heatmap of the corresponding correlation submatrix was also generated to complement the analysis.

## Results

### Differential Expression of CTAs

Among patients with available clinical data, the cohort of 156 GAC individuals was predominantly aged 60–69 years (20.5%) and male (47.4%). Most patients presented with stage III disease according to the TNM system (28.2%). Within the T, N, and M categories, the majority were classified as T4 (33.3%) and N3 (26.9%), while distant metastasis (M1) was observed in 19.2% of cases. Regarding Lauren classification, 48.1% were of the intestinal type and 22.4% of the diffuse type. According to TCGA classification, the most frequent subtypes were CIN (17.3%) and MSI (5.1%), although information was unavailable for 70% of cases. Most tumors were located in non-cardia regions (60.9%) (Table 1).

**Table 1.**
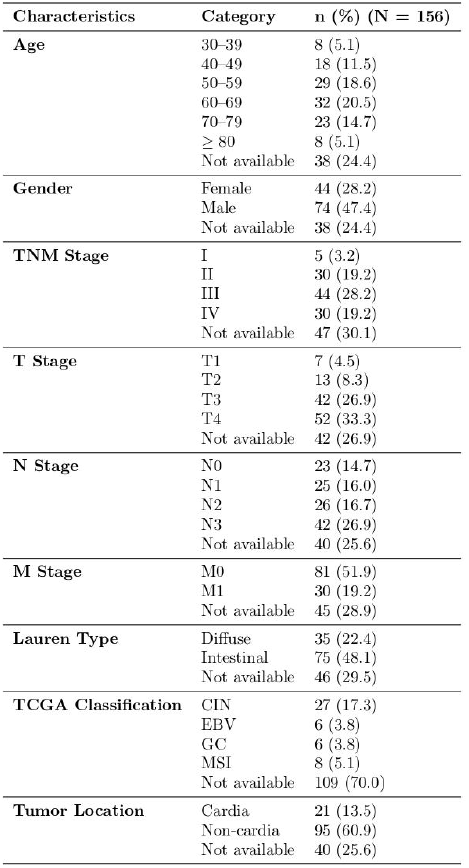
Clinicopatholpgical characteristics of patients with gastric adenocarcinoma.

The flowchart in Figure 1 illustrates the analytical workflow. The study included 362 samples, divided into GAC (156), PTT (186), and MP (20) tissues. Gene expression profiles of tumor, PTT, and MP samples were compared, resulting in a total of 541 DEGs identified based on the cohort criteria described above. Of these, 49 (9.06%) belonged to the previously defined CTA gene list. Differential expression analysis using the DESeq2 package in R across the different tissue samples is presented in Figure 2.

**Figure 1.**
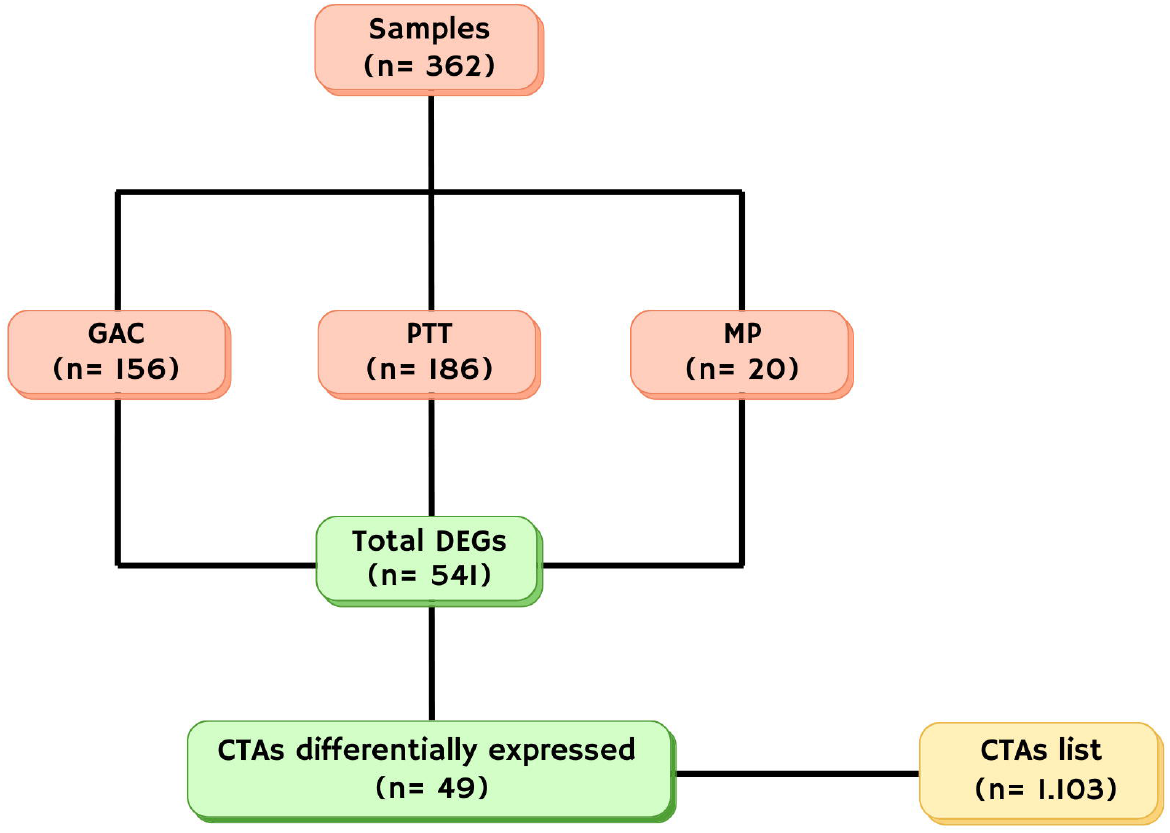
Flowchart of the analysis. Distribution of samples (n = 362) among the GAC, PTT, and MP groups, identification of DEGs, and selection of CTAs.

**Figure 2.**
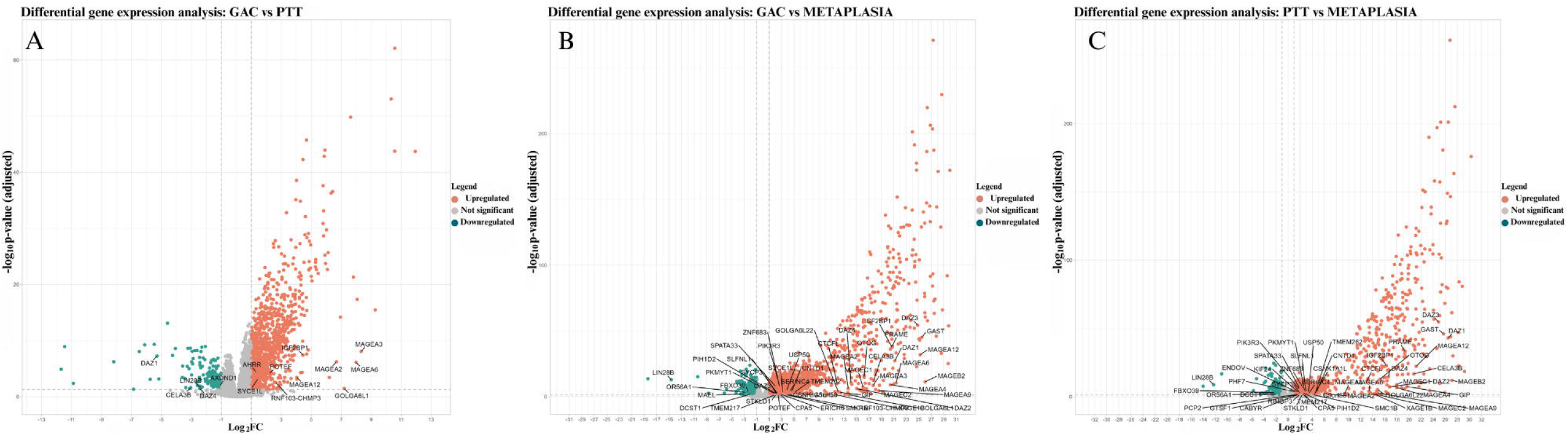
Volcano plot of DEGs between groups. (A) DEGs between GAC vs PTT. (B) DEGs between GAC and MP. (C) DEGs between PTT vs MP. The X-axis represents the L2FC of expression variation and the Y-axis corresponds to the −log10 of the adjusted p-value. Genes in red are upregulated, in green are downregulated, and in grey are not significant. Highlighted genes were selected for statistical and/or biological relevance.

Comparison between tumor and peritumoral tissues identified 14 differentially expressed CTA genes, of which 71.43% (10/14) were upregulated and 28.57% (4/14) were downregulated (Figure 2A). Comparing tumor and metaplastic tissues revealed 47 DEGs, with 91.49% (43/47) upregulated and 8.51% (4/47) downregulated (Figure 2B). CTA gene expression was also compared between peritumoral and metaplastic tissues, yielding 39 DEGs, comprising 92.31% (36/39) upregulated genes and 7.69% (3/39) downregulated genes (Figure 2C).

Among the 486 upregulated DEGs in the comparison between GAC and PTT, four CTAs were identified among the 15 most highly expressed genes in tumors (Table 2). Notably, MAGEA3 ranked 6th with an L2FC > 8, followed by MAGEA6 in 8th place (L2FC > 7). GOLGA6L1 and MAGEA2 were ranked 11th and 13th, respectively, with L2FC values >7 and >6.

**Table 2.** Top fifteen upregulated DEGs in GAC compared with PTT.

| Gene | L2FC | adjusted p-value | Category |
| --- | --- | --- | --- |
| ZIC1 | 11,9 | 0 | Others |
| STRA6 | 10,5 | 0 | Others |
| FOXC2 | 10,5 | 0 | Others |
| ZIC2 | 10,3 | 0 | Others |
| ITIH2 | 9,2 | 0 | Others |
| MAGEA3 | 8,3 | $8.45 \times 10^{-9}$ | CTA |
| SERPINB3 | 8 | 0 | Others |
| MAGEA6 | 7,9 | $7.33 \times 10^{-7}$ | CTA |
| CRABP1 | 7,8 | 0 | Others |
| FIBIN | 7,6 | 0 | Others |
| GOLGA6L1 | 7,2 | 0,03 | CTA |
| FBN3 | 6,9 | 0 | Others |
| MAGEA2 | 6,6 | $5.99 \times 10^{-7}$ | CTA |
| SFRP2 | 6,4 | 0 | Others |
| GAP43 | 6,3 | 0 | Others |

### Association of CTAs with Clinical Variables

Gene expression analysis revealed distinct associations with clinically relevant variables in GAC. Initially, normalized gene expression was used to assess the relationship between DEGs and variables of interest using the DESeq2 package in R. This approach identified 30 genes significantly associated with various clinical parameters, including TNM staging, T, N, and M categories, TCGA classification, Lauren classification, therapeutic management, and sex (Supplementary Figure 1). Additionally, paired contrast analyses between clinical variables were also performed (Figure 3).

**Figure 3.**
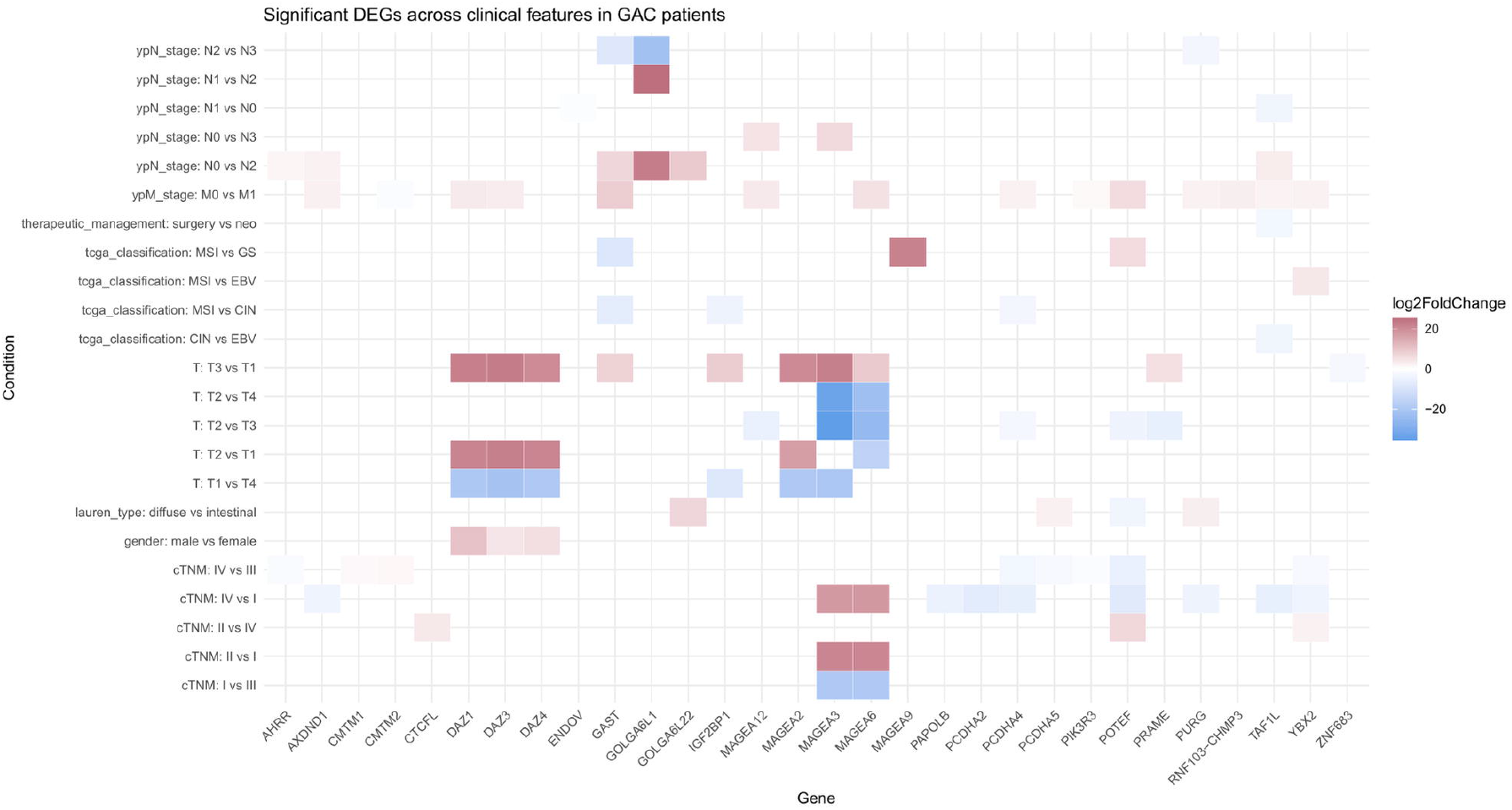
Heatmap of association between CTA expression and clinical variables in patients with GAC, obtained by differential expression analysis with DESeq2. Comparisons were performed in pairs between groups. The Y-axis represents the comparison in pairs of clinical subgroups (TNM staging, TCGA classification, Lauren classification, gender, and therapeutic management), and the X-axis represents CTAs genes. Each cell represents the L2FC value of a gene in the contrast between two groups, indicating in which of them the gene is most expressed.

Significant associations with the TNM system were observed for POTEF (FDR = 4.41×10^−6^ – 0.00816), MAGEA6 (FDR = 0.000395 – 0.00138), MAGEA3 (FDR = 0.00158 – 0.00461), and CTCFL (FDR = 0.0335). Additional candidates, including PIK3R3, CMTM1, TAF1L, YBX2, AHRR, CMTM2, PAPOLB, PURG, PCDHA2/4/5, and AXDND1, also showed significant differences in expression across tumor stages.

Regarding tumor invasion depth (T category), members of the MAGE family (MAGEA2, MAGEA3, MAGEA6, MAGEA12), POTEF, PRAME, IGF2BP1, DAZ1/3/4, PCDHA4, GAST, and ZNF683 were differentially expressed (FDR < 0.05). For the ypN category of staging, MAGEA3, MAGEA12, TAF1L, GAST, AHRR, GOLGA6L1, GOLGA6L22, PURG, AXDND1, and ENDOV were identified as significant candidates associated with lymph node involvement after treatment.

Analysis of the ypM category revealed strong associations for GAST (FDR = 3.16×10^−8^), POTEF (FDR = 1.42×10^−7^), PIK3R3 (FDR = 0.000557), YBX2 (FDR = 0.00123), PCDHA4 (FDR = 0.00248), and MAGE genes (MAGEA6, MAGEA12), emphasizing the role of these antigens in tumor dissemination. PURG, TAF1L, CMTM2, RNF103–CHMP3 and DAZ3 were also identified as significant.

MAGEA9 (FDR = 0.00695) and YBX2 (FDR = 0.00314) exhibited highly significant differences in expression among the TCGA molecular subtypes. Similarly, POTEF, IGF2BP1, GAST, and TAF1L also showed significant expression. Regarding Lauren classification, PURG (FDR = 0.000634), POTEF (FDR = 0.00824), GOLGA6L22 (FDR = 0.0189), and PCDHA5 (FDR = 0.0466) displayed significant differences in expression between diffuse and intestinal subtypes, indicating a distinct molecular profile between histological subtypes. In terms of therapeutic management (surgery vs. neoadjuvant therapy), TAF1L was the only gene showing a significant association (FDR = 0.00602). Finally, analysis of sex-related differences revealed strong associations for members of the DAZ family, including DAZ1 (FDR = 2.92×10^−13^), DAZ3 (FDR = 0.0452), and DAZ4 (FDR = 0.00138), reflecting sex-linked expression patterns.

As a complementary approach, classical non-parametric statistical tests were applied, including Kruskal–Wallis for multiple-group comparisons and Wilcoxon tests for pairwise comparisons, on the 30 genes identified as statistically significant in the DESeq2 analysis. Results showed that eight CTA genes maintained significant associations across different clinical subgroups (Figure 4).

**Figure 4.**
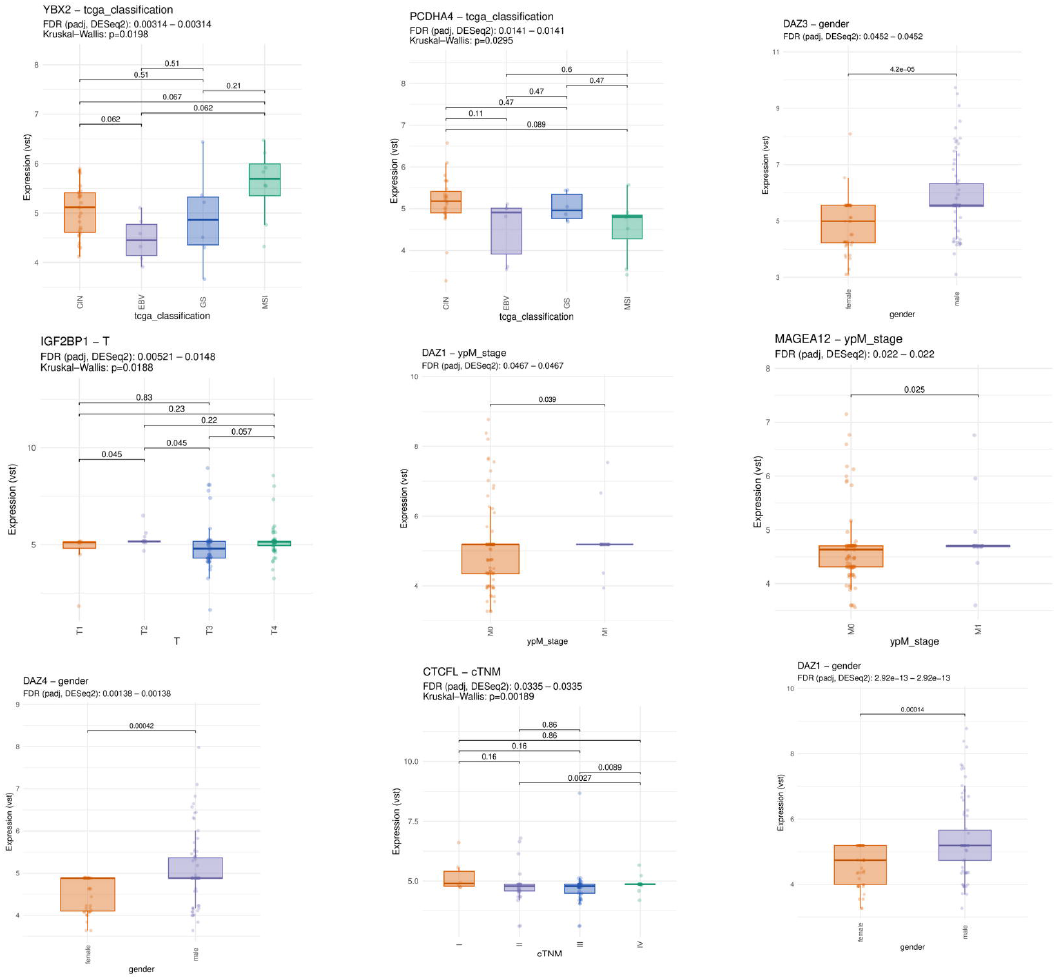
Boxplot of CTA expression in association with clinical variables of patients with GAC. Each graph shows the distribution of normalised expression of a gene in different clinical categories, including TNM staging, T category, gender, and M category. The FDR values shown were obtained by differential expression analysis in DESeq2. Additional comparisons between groups for each clinical variable were performed using classical statistical tests (p-values displayed above the bars), including paired contrasts between levels of the same variable.

The gene CTCFL exhibited significantly increased expression (p = 0.00189), and pairwise comparisons revealed higher expression of this gene in stage IV, both between stages II and IV (p = 0.0027) and between III and IV (p = 0.0089), suggesting a potential role for CTCFL in tumor progression. Similarly, IGF2BP1 showed a potential positive association with tumor T category (p = 0.0188), and in pairwise comparisons, increased expression was observed in patients classified as T2 in both T1 vs. T2 and T2 vs. T3 comparisons (p = 0.045 for both).

DAZ1 displayed statistically significant associations with sex and ypM stage, with higher expression in male patients (p = 0.00014) and in the direct comparison of M0 vs. M1 (p = 0.039). Greater variability and higher expression levels were observed in non-metastatic tumors (M0), whereas metastatic tumors (M1) exhibited more homogeneous and lower expression. In addition, DAZ4 and DAZ3 were more highly expressed in males, with p-values of 0.00042 and 4.2×10^−5^, respectively.

The expression of MAGEA12 was notably higher in M1 tumors when comparing the M category groups (p = 0.025). In the post-treatment ypM stage, MAGEA12 expression differed significantly between M0 and M1 patients (p = 0.025), with M0 patients showing greater heterogeneity, including higher expression levels in some cases, while M1 patients exhibited more uniform expression.

The genes PCDHA4 and YBX2 showed significant differential expression among the TCGA molecular subtypes (p = 0.0295 and p = 0.0198, respectively). For PCDHA4, expression was higher in CIN tumors compared to MSI, although pairwise comparisons were only suggestive. In contrast, YBX2 exhibited increased expression in the MSI subtype compared to the other groups (CIN, EBV, and GS), with borderline significance in multiple-group comparisons.

### Survival Analysis

Kaplan–Meier curve analysis revealed that high expression of several genes was associated with worse prognosis in GAC. These include DAZ3, DAZ1, GAST, CT45A5, MAGEA2, MAGEC1, MAGEC2, POTEF, LIN28B, PCDHA4, OTOG, GOLGA6L22,IGF2BP1, and CTCFL, all with hazard ratios (HR) above 1, indicating a significantly increased risk of death in patients with elevated expression (Figure 5). Notably, IGF2BP1 (HR = 2.86, p = 0.004), CTCFL (HR = 2.77, p = 0.0039), CT45A5 (HR = 2.81, p = 0.0033), and LIN28B (HR = 2.72, p = 0.0045) showed the highest relative risks, underscoring their potential role as markers of tumor aggressiveness. In contrast, SYCE1L (HR = 0.56, p = 0.040), PIK3R3 (HR = 0.50, p = 0.012), and ZNF683 (HR = 0.56, p = 0.045) were associated with improved overall survival.

**Figure 5.**
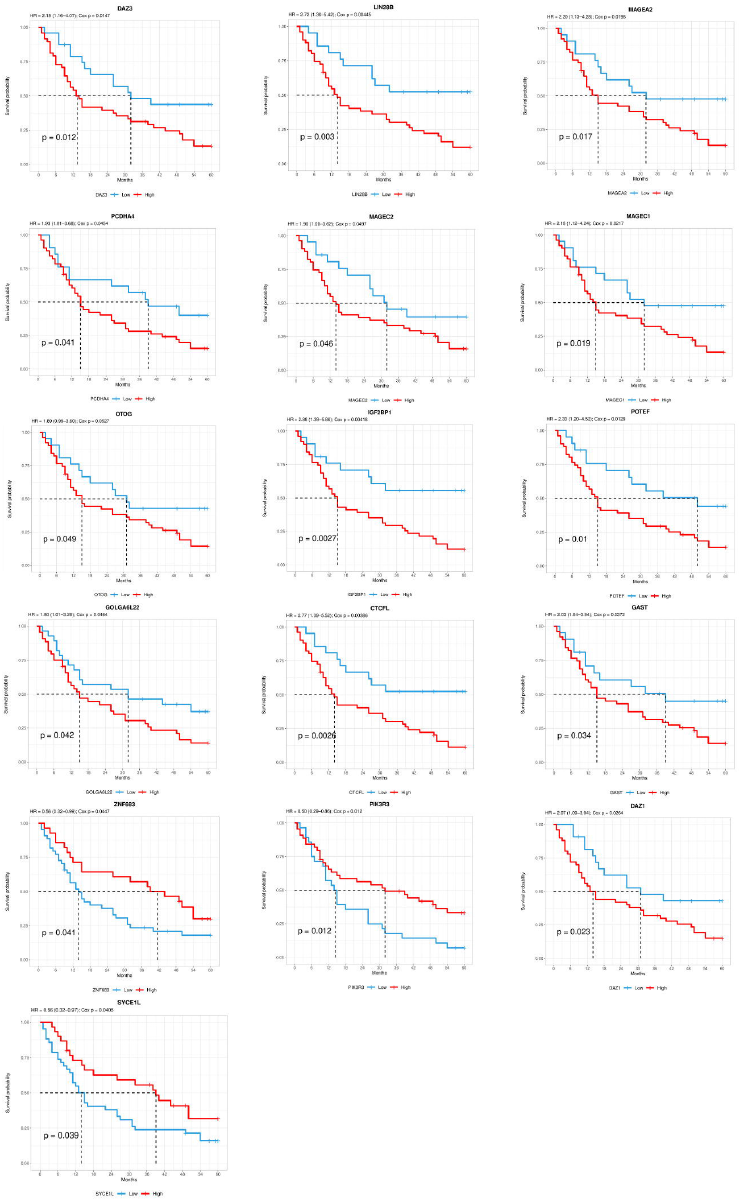
Kaplan-Meier curves for overall survival analysis in patients with GAC, according to gene expression levels. The curve represent the probability of survival over time (in months) comparing patients with high expression (red line) and low expression (blue line) of the genes: DAZI, DAZ3, MAGEA2, LIN28B, PCDHA4, MAOECl-2, OTOG, IGF2BPI, POTEF, GOLGA6L22, CTCFL, GAST, ZNF683, PIK3R3, and SYCEI L.

### Co-expression Network Analysis

The co-expression network constructed from the differentially expressed CTAs revealed five main clusters (Figure 6). Cluster 1 (dark blue) grouped members of the DAZ family (DAZ1–DAZ4), which were strongly correlated with each other, with DAZ1, DAZ3, and DAZ4 emerging as the main hub genes. This cluster also showed moderate positive correlations with AADAC, FBP2, and OVCH2.

**Fig. 6.**
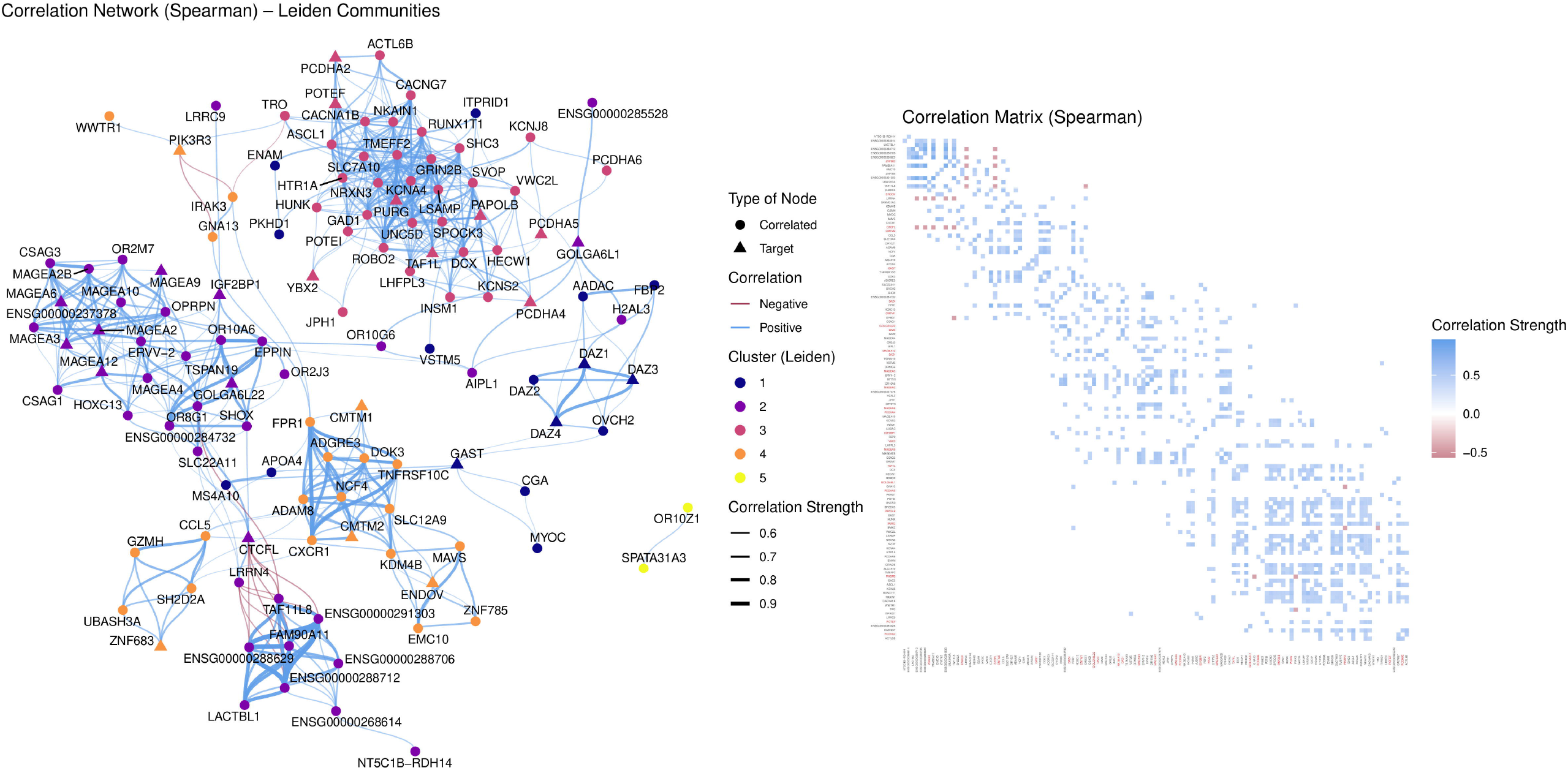

Cluster 2 (purple) comprised members of the MAGEA family (MAGEA2/2B/3/4/6/9/10/12), which exhibited strong correlations among themselves, along with additional connections to OPRPN, OR2M7, and CSAG3, forming a highly interconnected germline–tumor core. Key hub genes in this cluster included MAGEA2, MAGEA3, MAGEA6, MAGEA9, MAGEA12, IGF2BP1, CTCFL, GOLGA6L1, and GOLGA6L22, with the latter two showing weaker correlations with the rest, possibly indicating a more peripheral functional role.

Cluster 3 (pink) encompassed YBX2, POTEF, members of the PCDHA family (PCDHA2, PCDHA4, PCDHA5), TAF1L, PAPOLB, and PURG, all of which were tightly interconnected. These genes, together with their respective partners, appear to be associated with cell adhesion and transcriptional regulation processes.

Cluster 4 (orange) included genes related to immune and inflammatory responses, such as CXCR1, CCL5, GZMH, ADAM8, TNFRSF10C, UBASH3A, and IRAK3. Notably, this cluster showed weak but negative correlations between genes such as IRAK3, GNA13, and PIK3R3 and regulatory CTAs (e.g., MAGEA, CTCFL, IGF2BP1), suggesting a potential functional antagonism between proliferative and inflammatory axes. Relevant target genes in this context included PIK3R3, CMTM1, CMTM2, ENDOV, and ZNF683.

Finally, Cluster 5 (yellow) displayed low connectivity, containing only SPATA31A3 and OR10Z1, considered target genes of a possible accessory module whose functional relevance remains to be elucidated.

## Discussion

Gastric cancer (GC) remains a major public health issue, and recent estimates indicate that global mortality from this disease will remain substantial in the coming years [28]. In this context, the identification of novel therapeutic targets and biomarkers is essential to improve risk stratification and expand available treatment options.

The ideal antigen for cancer therapies should be tumor-specific, highly immunogenic, broadly expressed in tumor cells, shared among patients, and indispensable for tumor survival [29]. Cancer-testis antigens (CTAs) meet most of these criteria, making them attractive candidates for immunotherapy [30]. However, for this approach to be translationally useful, it is necessary to comprehensively characterize their expression patterns across different stages and clinical contexts of gastric cancer (GC).

Although recent studies have investigated the expression of these genes in gastric cancer (GC), the use of restricted CTA lists for analysis represents a limiting factor. Additionally, analyses comparing tumor tissue with peritumoral tissue (PTT) and premalignant metaplastic tissue (MP) may provide complementary information. Furthermore, assessing gene expression in association with clinically relevant prognostic variables can offer new insights.

In this study, we identified 49 differentially expressed CTAs across the analyzed contrasts. Between gastric cancer (GC) and peritumoral tissue (PTT), only 14 DEGs were observed, whereas the comparison between GC and metaplastic tissue (MP) revealed 47 DEGs. Notably, a large proportion of genes upregulated in the tumor compared to MP (23 genes) were also positively regulated in PTT relative to MP. These findings suggest that peritumoral tissue is more similar to tumor tissue than MP, which can be explained by the field cancerization process, in which regions adjacent to the tumor already harbor genetic and/or epigenetic alterations that render them biologically closer to the tumor and predisposed to the development of new lesions or local recurrence [31]. In contrast, the comparison between PTT and MP resulted in 39 DEGs, highlighting a high degree of divergence between these two tissues.

In the comparison between GC and PTT, among the 486 upregulated DEGs, four CTAs stood out among the 15 most abundant: MAGEA3, MAGEA6, GOLGA6L1, and MAGEA2. In addition, IGF2BP1, MAGEA12, RNF103-CHMP3, and POTEF were also positively regulated in the tumor. Notably, all these genes maintained the same expression pattern in the comparison between GC and MP, suggesting their potential as malignancy markers and therapeutic targets. However, MAGEA3, MAGEA6, MAGEA2, IGF2BP1, and MAGEA12 were also upregulated in PTT relative to MP, indicating that, although not tumor-exclusive, they may serve as risk biomarkers.

Conversely, LIN28B, FBXO39, and OR56A1 were downregulated in GC and PTT compared to MP. Notably, LIN28B was also negatively regulated in the tumor relative to PTT. This pattern suggests that these genes are more active in tissues with premalignant alterations and become progressively silenced during tumorigenic transformation. Such an expression profile may render them potential progression biomarkers, indicating the transition from metaplasia to tumor. These genes could also be valuable for the early detection of tissues adjacent to malignant transformation during gastric carcinogenesis.

CTCFL expression was significantly associated with TNM stage and patient survival. Although elevated even at early stages, expression was more pronounced in stage IV, suggesting a role in tumor progression. As a transcription factor, CTCFL is involved in the deregulation of oncogenic promoters, including c-MYC, hTERT, p53, INK/ARF, MDM2, PLK, and PIM [32]. In melanoma, CTCFL has been shown to promote the transition from an intermediate phenotype to an invasive mesenchymal state, associated with increased chromatin accessibility and oncogene instability [33,34].

This profile is consistent with our clinical findings, in which patients with high CTCFL expression exhibited reduced survival and a 2.77-fold increased risk of death. Additionally, positive regulation was observed in both GC and PTT compared to MP. Collectively, these results indicate that CTCFL may serve as a biomarker of advanced stage and prognosis in gastric cancer, as well as a promising therapeutic target to impede tumor progression.

IGF2BP1 functions as a post-transcriptional regulator in cancer cells, stabilizing pro-oncogenic mRNAs and contributing to proliferation, growth, invasion, chemoresistance, metastasis, and reduced overall survival. Its involvement has been described in various cancer types, including esophageal, lung, melanoma, glioblastoma, liver, and gastrointestinal tumors [35,36]. Furthermore, it has been shown that the lncRNA GHET1, which is highly expressed in GC, physically interacts with IGF2BP1, enhancing its binding to c-MYC mRNA and promoting its stabilization, resulting in increased oncogenic activity [35,37].

In our study, IGF2BP1 expression was associated with T2 stage within the T category of the TNM system, suggesting its involvement in early events of muscular layer invasion and the potential transition from a localized tumor to an invasive phenotype. Furthermore, overexpression demonstrated prognostic value, as patients with high IGF2BP1 levels exhibited a 2.85-fold increased risk of adverse clinical outcomes.

The high expression of DAZ1, DAZ3, and DAZ4 in men is consistent with the localization of the DAZ gene cluster on the Y chromosome and reflects their role in spermatogenesis. However, high expression of DAZ3 and DAZ1 was also associated with poorer SG. Comparative analysis revealed that DAZ4 and DAZ1 are downregulated in the tumor relative to PTT, whereas DAZ3 is upregulated in both tumor and PTT compared to MP. In breast cancer cell lines, high DAZ1 expression has been observed [38]; however, the underlying mechanisms remain unclear, and further investigation is required in the context of our study.

The MAGEA12 gene also showed significantly increased expression in cases classified as M1, indicating an association with the presence of distant metastasis. Furthermore, this gene was positively regulated in the tumor compared to both MP and PTT. Similarly, MAGEA12 was upregulated in PTT relative to metaplasia. These findings suggest that MAGEA12 activation is not limited to the tumor itself but may extend to adjacent tissue, potentially contributing to a more aggressive phenotype and metastatic capacity. MAGEA12 overexpression has also been reported in gastric and cutaneous squamous cell carcinoma, breast cancer, and hepatocellular carcinoma [39,40,41,42].

High expression of ZNF683, PIK3R3, and SYCE1L appears to be associated with a more favorable profile, potentially linked to the modulation of immune response pathways. These genes may serve as useful prognostic biomarkers, aiding in patient stratification and the identification of subgroups with better clinical outcomes.

Correlation analysis revealed five distinct clusters among the differentially expressed CTAs, each organized around central target genes. Cluster 1, comprising DAZ family genes, indicates reactivation of germline programs in cancer [43, 44, 45]. Cluster 2, formed by MAGEA genes, showed strong correlations with other regulatory CTAs, such as IGF2BP1 and CTCFL, suggesting a central role in proliferation and immune evasion [46, 47]. Cluster 3 included genes associated with cell adhesion and transcriptional regulation, including YBX2, POTEF, and members of the PCDHA family, also serving as target genes [48, 49]. Cluster 4 grouped genes related to immune response, such as PIK3R3 and ZNF683, exhibiting negative correlations with proliferative CTAs, suggesting potential antagonism between tumorigenic and inflammatory pathways [50, 51, 52]. Finally, Cluster 5 displayed low connectivity, with SPATA31A3 and OR10Z1 as target genes potentially linked to accessory functions. These findings highlight the functional organization of CTAs into correlation clusters, with potential implications for tumor regulation and immune response.

The integrated analysis of cancer-testis antigens (CTAs) expression in GAC identified genes with strong clinical and prognostic relevance. Genes such as IGF2BP1, CTCFL, and MAGEA12 were associated with advanced stages, presence of metastasis, and poorer overall survival, whereas PIK3R3, SYCE1L, and ZNF683 exhibited a protective effect. Coexpression network analysis revealed germline, regulatory, and immune modules, indicating functional antagonism between proliferative and immune response pathways. Altogether, these findings underscore the potential of CTAs as biomarkers and therapeutic targets for immunotherapy strategies in GC.

Although this study provides a comprehensive characterization of CTAs in GAC, several limitations should be acknowledged. The use of retrospective RNA-Seq data without experimental validation limits the generalizability of the findings, and missing clinical information restricts the robustness of the observed associations. Furthermore, the lack of consensus regarding the classification of certain genes as CTAs may introduce potential biases. Coexpression analyses, while informative, do not establish causal relationships and therefore require additional functional validation. Finally, expression was assessed only at the transcriptional level, without consideration of protein-level data. These limitations highlight the need for future studies integrating experimental validation and multi-omics approaches to consolidate and expand the clinical utility of the identified biomarkers.

## Acknowledgments

The authors thank Fundação Amazônia de Amparo a Estudos e Pesquisas (FAPESPA) and Conselho Nacional de Desenvolvimento Científico e Tecnológico (CNPq) for funding this study. We also acknowledge the Oncology Research Center (NPO) at the Federal University of Pará (UFPA) for technical support and for providing the resources necessary to carry out this research.

