## Supplementary figure 1 for "Cancer-Testis Antigens as Clinical and Prognostic Biomarkers in Gastric Adenocarcinoma: Integration of Differential Expression, Clinical Associations, Survival, and Co-Expression Networks"

### Significant genes for the variable: cTNM

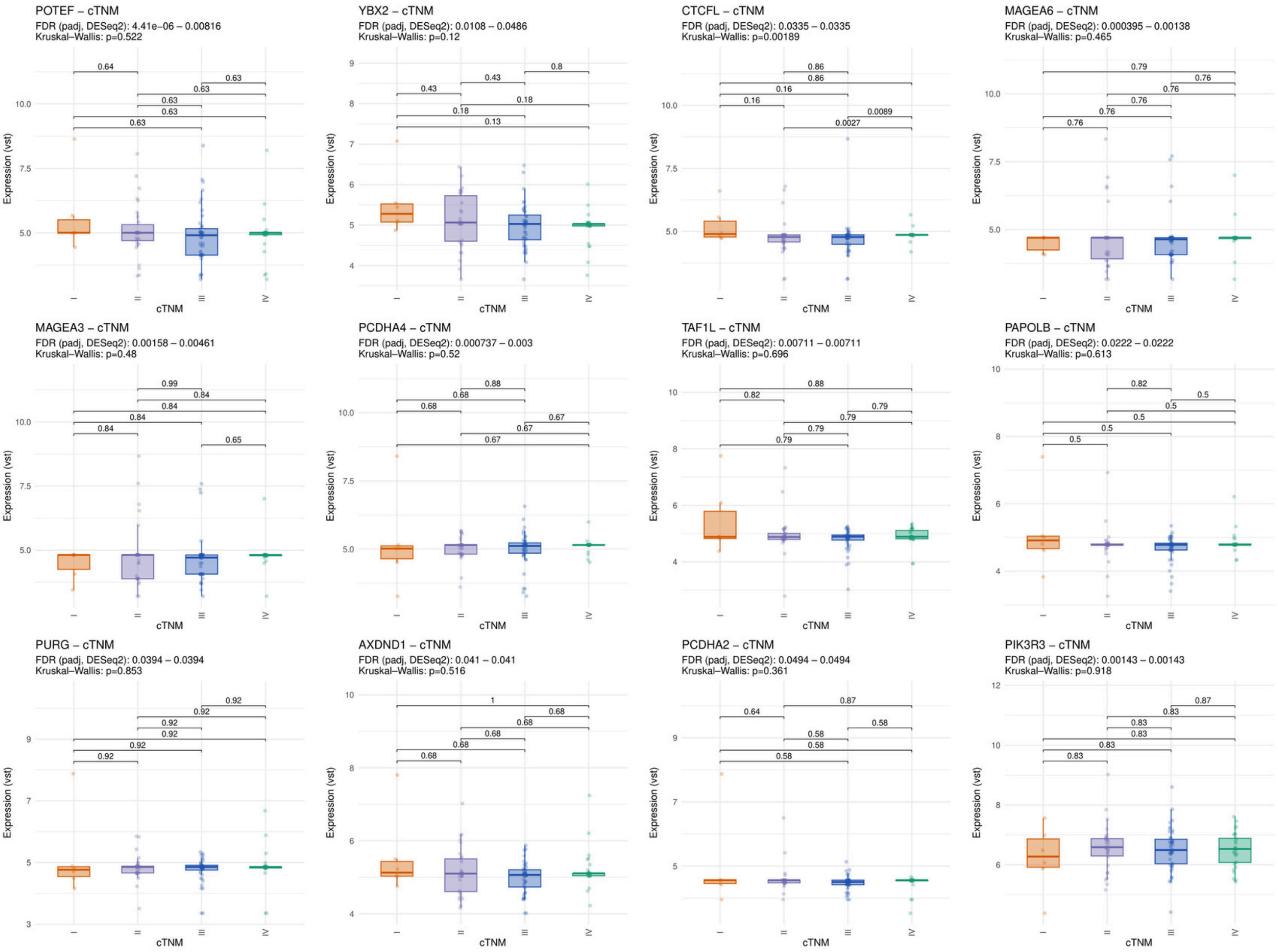

### Significant genes for the variable: cTNM

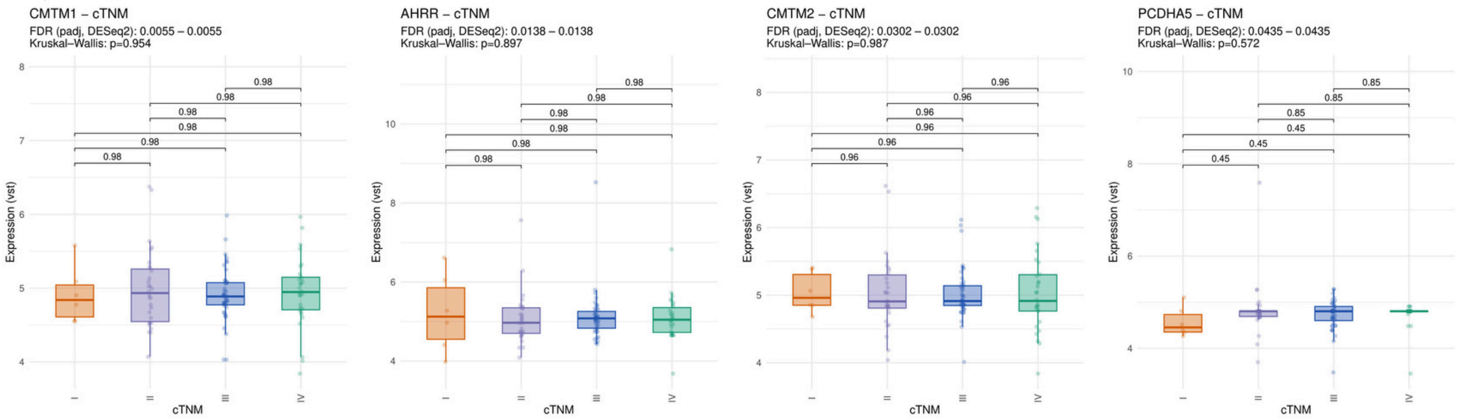

Significant genes for the variable: ypT_stage

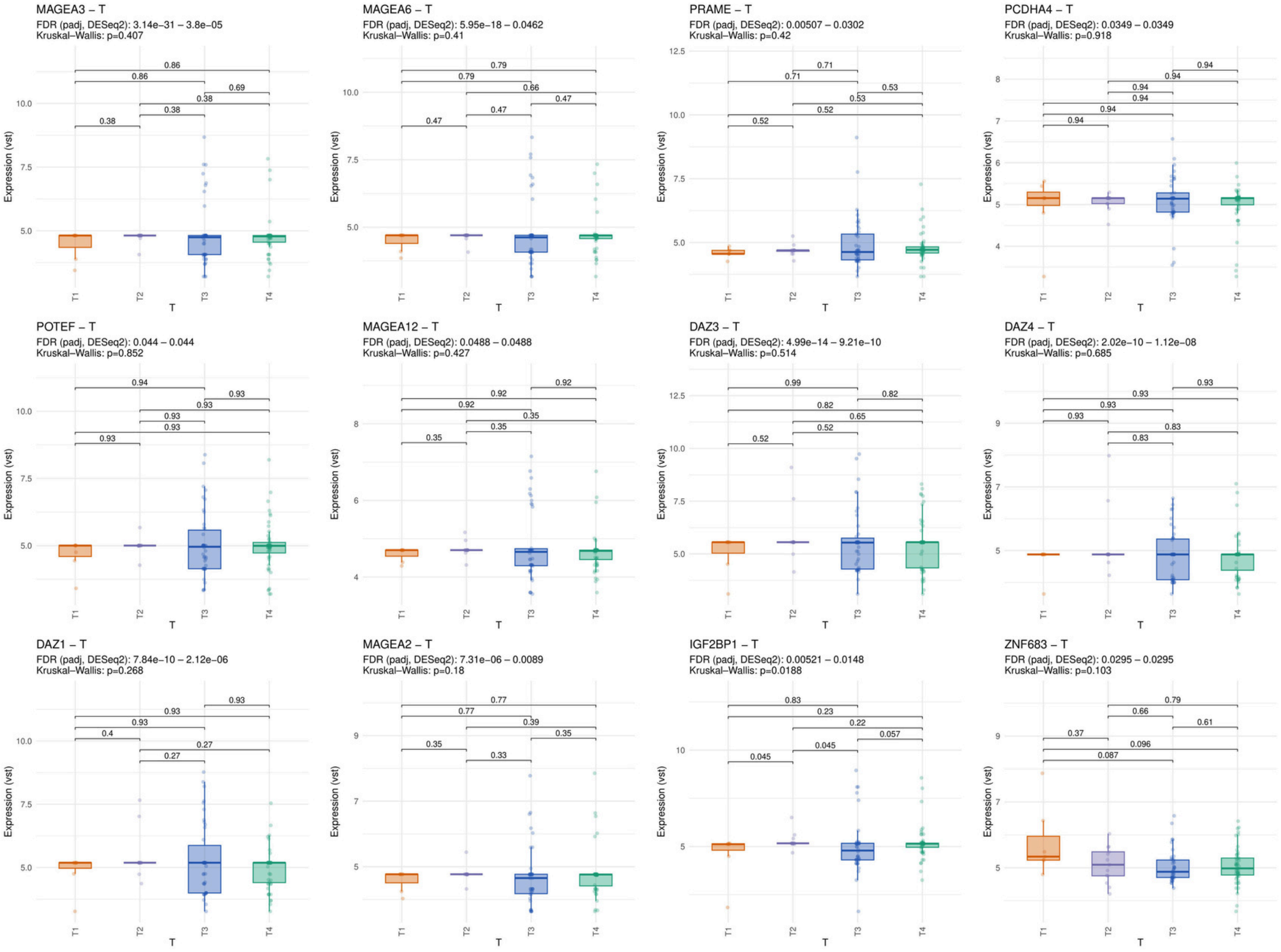

### Significant genes for the variable: ypT_stage

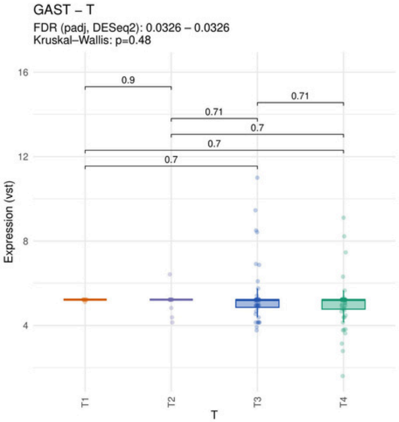

### Significant genes for the variable: ypN_stage

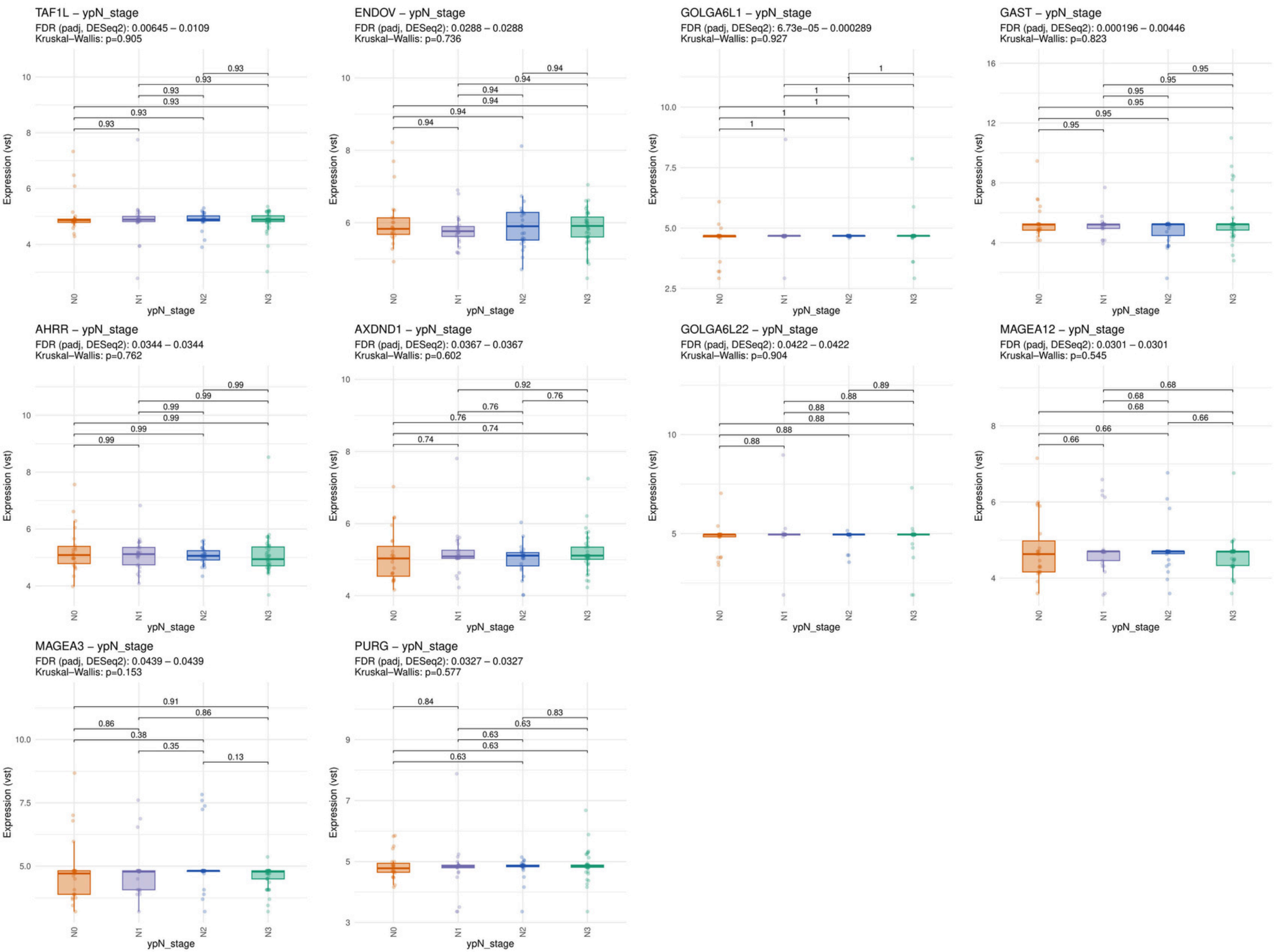

Significant genes for the variable: ypM_stage

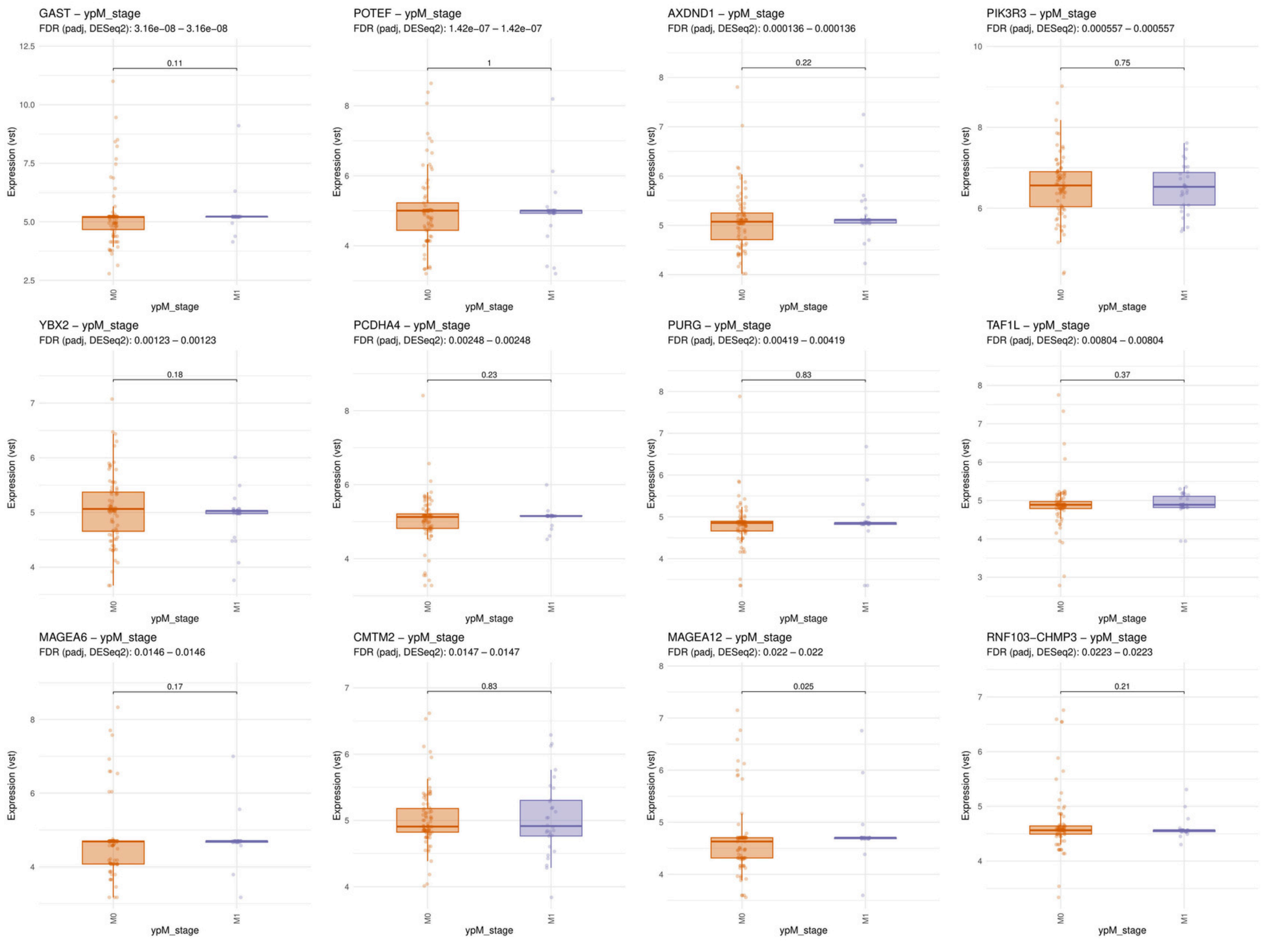

### Significant genes for the variable: ypM_stage

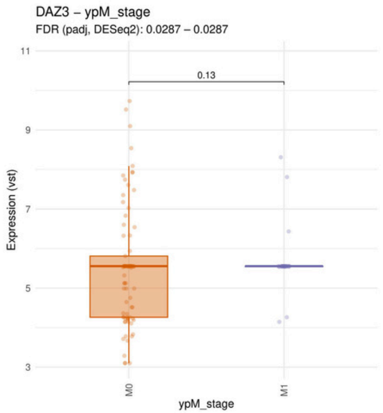


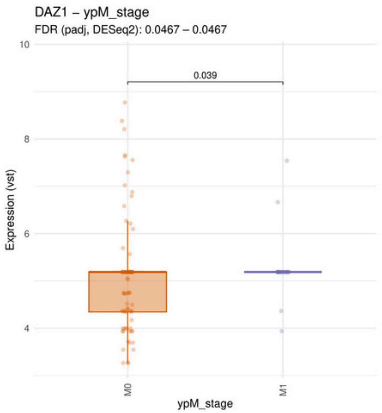

#### Significant genes for the variable: TCGA_classification

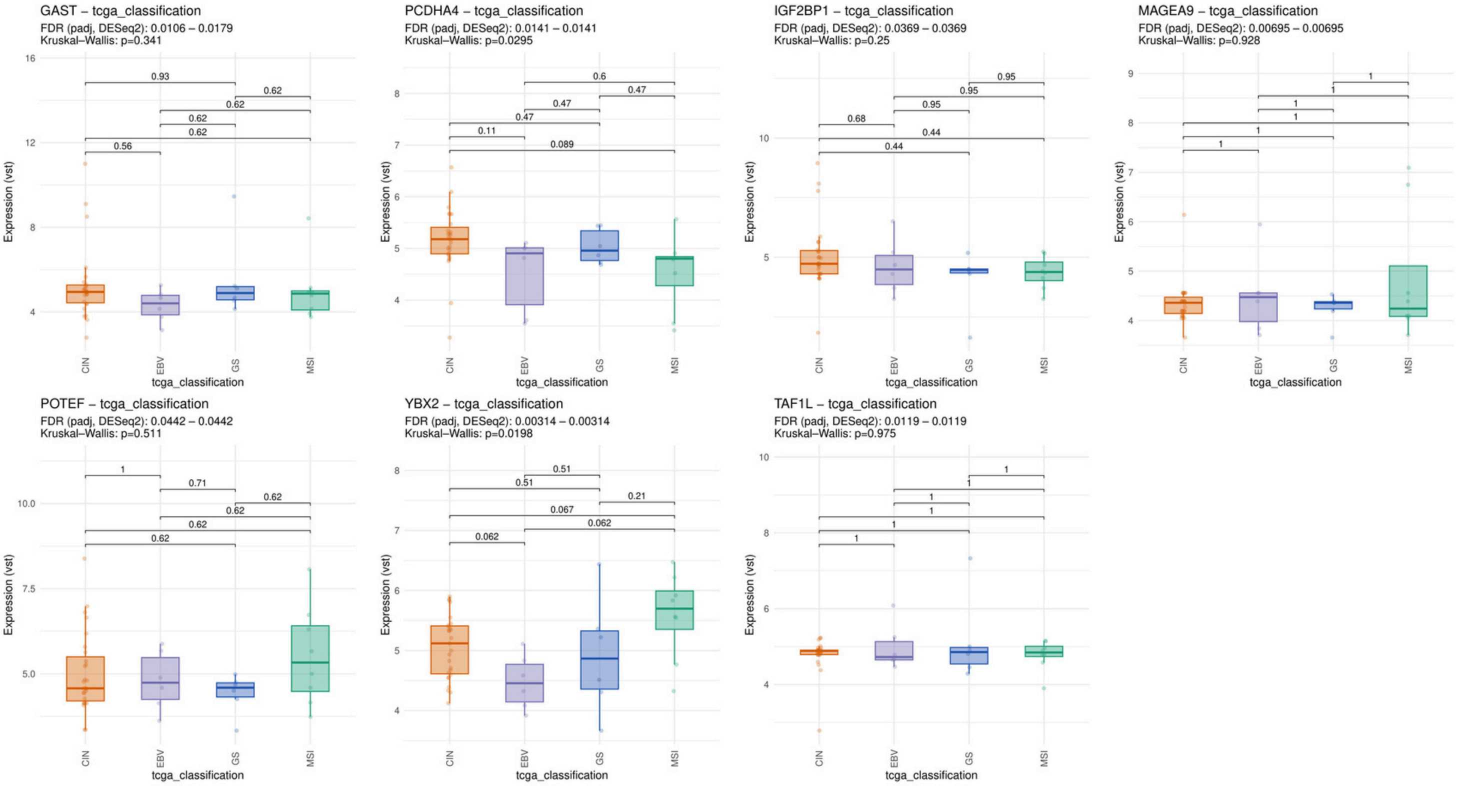

Significant genes for the variable: lauren_type

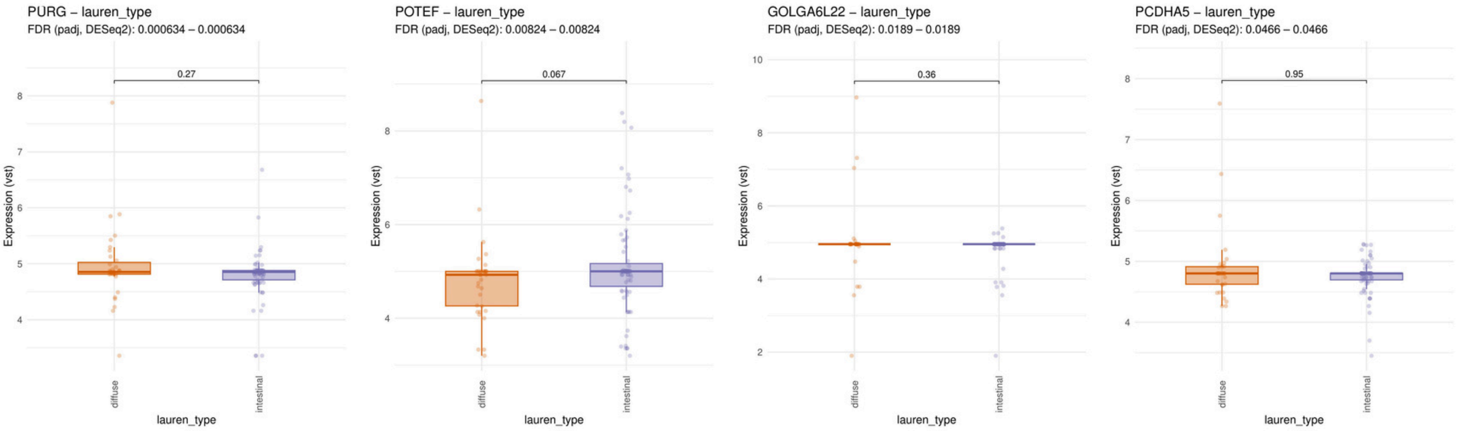

### Significant genes for the variable: therapeutic_management

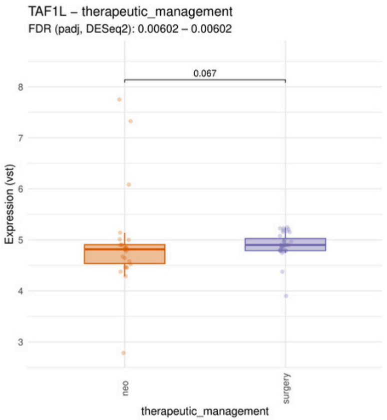

Significant genes for the variable: gender

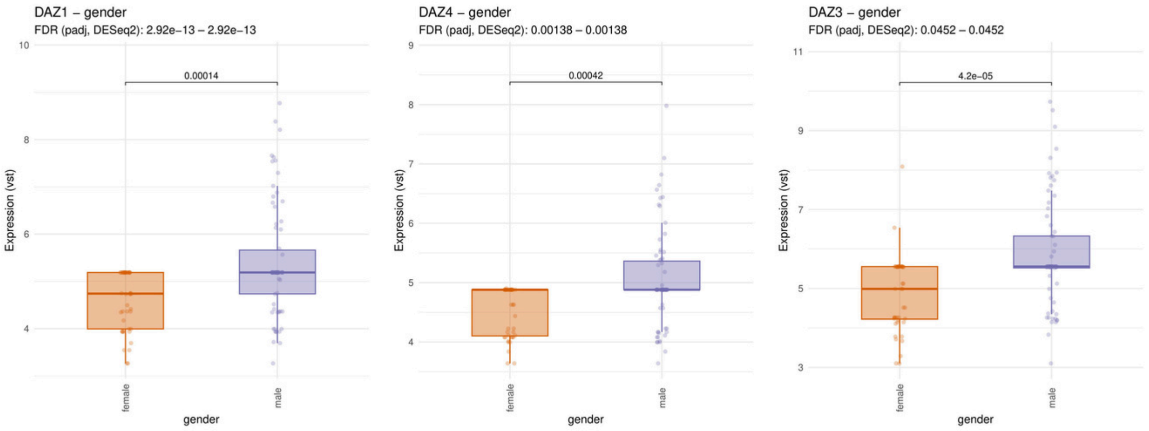
